# Light and noise pollution impacts specialist wildlife species disproportionately

**DOI:** 10.1101/2021.02.18.431905

**Authors:** Mark A. Ditmer, Clinton D. Francis, Jesse R. Barber, David C. Stoner, Brett M. Seymoure, Kurt M. Fristrup, Neil H. Carter

## Abstract

Global expansion of lighting and noise pollution alters how animals receive and interpret environmental cues. Yet we lack a cross-taxon understanding of how animal traits influence species vulnerability to this growing phenomenon. This knowledge is needed to improve the design and implementation of policies that mitigate or reduce sensory pollutants. We present results from an expert knowledge survey that quantified the relative influence of several ecological, anatomical, and physiological traits on the vulnerability of terrestrial vertebrates to elevated levels of anthropogenic lighting and noise. Our findings, based on 280 responses, highlight the increasing recognition among experts that sensory pollutants are important to consider in management and conservation decisions. Participant responses show mounting threats to species with narrow niches; especially habitat specialists, nocturnal species, and those with the greatest ability to differentiate environmental visual and auditory cues. Our results call attention to the threat specialist species face and provide a generalizable understanding of which species require additional considerations when developing conservation policies and mitigation strategies in a world altered by expanding sensory pollutant footprints. We provide a step-by-step example for translating these results to on-the-ground conservation planning using two species as case studies.

## Introduction

All organismal interactions with their environments are mediated by sensory inputs. Global growth in two sensory pollutants, anthropogenic lighting and noise (henceforth, “lighting” and “noise”), are fundamentally altering visual and auditory performance for many species (Dominoni et al. 2020). Light pollution and noise are pervasive, growing, and intensifying (Buxton et al. 2017; Kyba et al. 2017); altering sensory environments at a global scale. Importantly, all organismal interactions with their environments are mediated by sensory inputs, and these sensory pollutants both disrupt environmental cues and ecological processes near their source and extend far beyond the altered landcover (Barber, Crooks & Fristrup 2010; Kyba et al. 2015b). Approximately 80% of the IUCN’s Global Key Biodiversity Areas experience excess nightlight luminance (Garrett, Donald, & Gaston 2019) and 12% of the IUCN’s designated wilderness areas in North America experience anthropogenic noise above natural levels (Buxton et al. 2017).

Numerous studies assessing the impacts of lighting or noise provide examples of altered behaviors, and fitness costs have been documented (Longcore & Rich 2004; Francis & Barber 2013; Gaston et al. 2013). Evaluating these examples from an evolutionary perspective can reveal selective forces arising from novel stimuli (Swaddle et al. 2015; Hopkins et al. 2018) and identify the plausible taxonomic and ecological extents of similar effects. Noise reduces the ability to perceive acoustic signals while lighting affects visual perception. Both can fundamentally alter spatial orientation and create mismatched biological timings (Gaston et al. 2017). These sensory disturbances in turn create a myriad of behavioral alterations, affecting orientation and movement (Slabbekoorn & Bouton 2008; Cabrera-Cruz, Smolinsky, & Buler 2018), communication (Francis & Barber 2013), foraging and hunting efficiency (Bennie et al. 2015; Bunkley & Barber 2015; Mason, McClure, & Barber 2016), altered energy budgets (Read et al. 2014; Touzot et al. 2019) and predation risk (Francis & Barber 2013; Ditmer et al. 2020a), along with increased physiological stress (Rolland et al. 2012; Ouyang, Davies, & Dominoni 2018).

Most impacts of these sensory pollutants have been demonstrated with a relatively small number of species at local scales, primarily within North America and Europe. Studies involving terrestrial mammals are especially scarce (Shannon et al. 2016). Nonetheless, recent research has also shown that variation in these sensory pollutants better explains patterns of habitat selection than common ecological variables, such as landcover (Kleist et al. 2017), and better reflects the dynamic human footprint relative to other measurements (e.g., housing density; Ditmer et al. 2020b). Biological and ecological traits have frequently been linked with species’ vulnerability to environmental change and threat of extinction (Chown 2012; Foden et al. 2013), but with limited treatment in the context of vulnerabilities to lighting and noise (primarily avian species; see Francis 2015; Senzaki et al. 2020). Therefore, for species where scientific studies on the impacts of sensory pollutants are scarce or nonexistent, assessing relationships among relevant traits and taxa is most practical for forecasting species’ responses, and developing policies and conservation actions.

Here, synthesizing knowledge from experts around the world, we ranked the degree to which a range of ecological, anatomical, and physiological traits contribute to a species’ vulnerability to lighting and noise. Vulnerability is considered a function of exposure to a threat, sensitivity to the threat, and the corresponding adaptive capacity (McCarthy et al. 2001). Given the lack of published data on the subject across different taxa, we used an expert knowledge elicitation. This method has successfully been used to develop conservation policy (Martin et al. 2012), especially for subjects with incomplete scientific understanding (Foden et al. 2013), emerging threats (Klein et al. 2017), or resource limitations that preclude in depth studies (Carwardine et al. 2012; Gerber et al. 2018). Our survey did not ask questions regarding exposure to sensory pollutants, because levels may vary greatly within and among species and regions. Instead we followed the approach by Foden et al. (2013), who incorporated data from expert surveys to quantify the degree to which biological traits – reflecting sensitivity (i.e., the degree to which the survival, persistence, fitness, performance, or regeneration of a species is reliant on current night light and noise levels or characteristics) and adaptive capacity (i.e., the capacity of the species to persist in situ, shift to suitable microhabitats, or migrate to suitable regions [Dawson et al. 2011]) – influenced the threats of climate change. We collected independent responses from numerous experts, defined as sensory ecologists or individuals with ~3 or more years of study/experience with a vertebrate species/genera/taxa, that have diverse experiences with a variety of species across geographic and ecological regions. By leveraging the experience and ecological knowledge of these experts, our aim was to identify the traits that influence vulnerability to lighting and noise the most, such that any species can be preliminarily assessed for their likelihood to respond to these sensory pollutants as has been done for species vulnerability to climate change (Dawson et al. 2011).

## Materials and Methods

### Survey development and design

Using the methods of Foden et al. (2013), we designed an online survey (hosted at Qualtrics.com) to assess how and to what degree select biological traits contribute to anthropogenic light and noise vulnerability. We informed participants that the survey was only considering direct, negative impacts of lighting and noise on adult, terrestrial vertebrates. Traits incorporated into our survey were selected from recommendations developed at a 3-day workshop of experts in sensory ecology and animal physiology, where the effects of sensory pollutants and the mechanisms of disturbance were extensively discussed across a diverse range of taxa (Dominoni et al. 2020). We selected traits that were associated with increased extinction risk and allowed participants to complete the survey regardless of their primary species or taxa studied. As such, respondents were instructed not to consider idiosyncratic responses, but to focus on traits that are generally linked to increased vulnerability across vertebrate taxa. We provided definitions (and some examples) of each trait considered in the survey (**Table 1**), and we based our definition of vulnerability (provided to respondents) on the 2001 Assessment Report, in which vulnerability is a function of exposure, sensitivity, and adaptive capacity (McCarthy et al. 2001). We classified traits as either related to ecological, anatomical/physiological sensitivity, or adaptive capacity (**Table 1**). However, “use of migration” was the only trait classified as adaptive capacity, so we grouped it with the ecological sensitivity traits in the Results.

**Table 1.).**
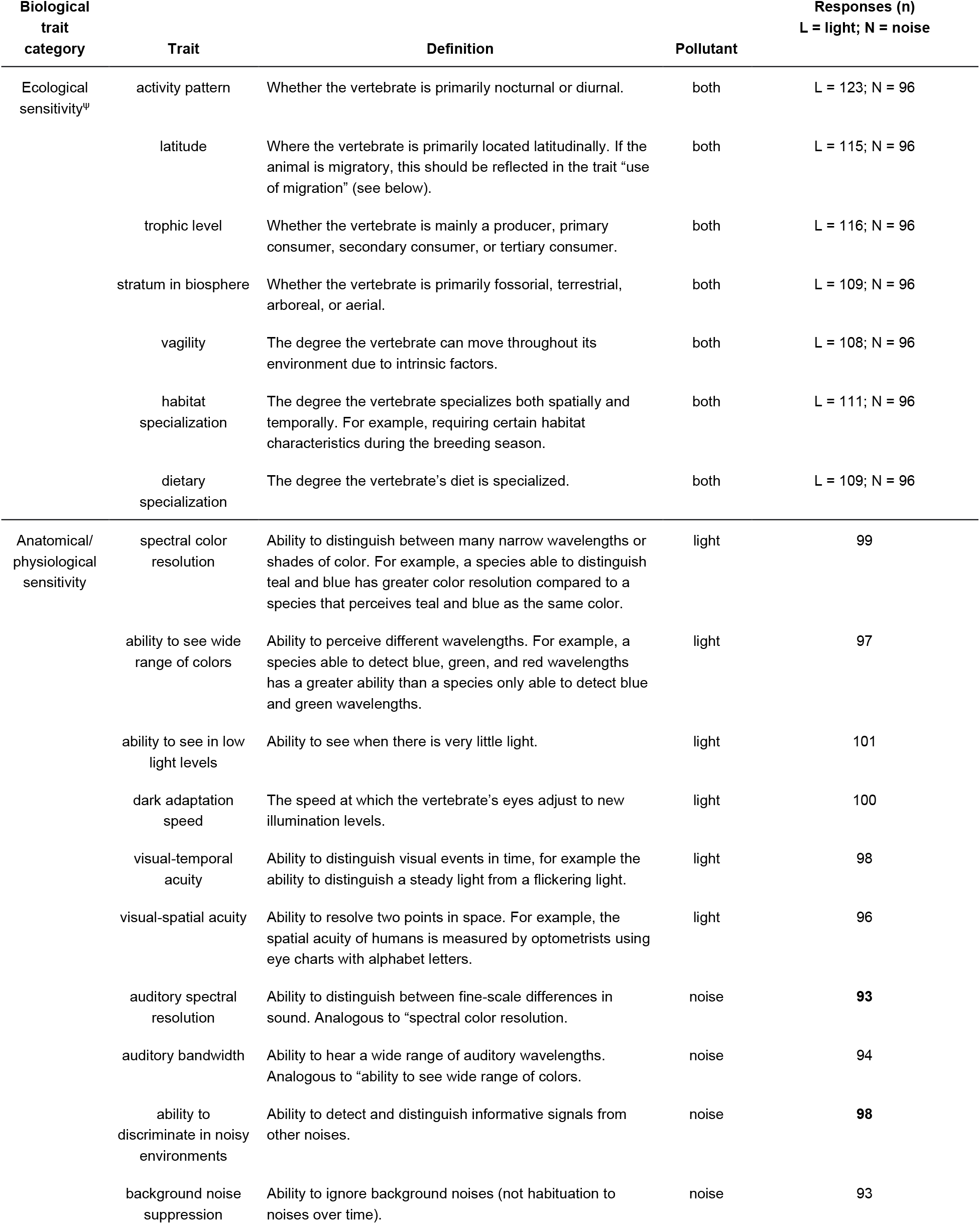
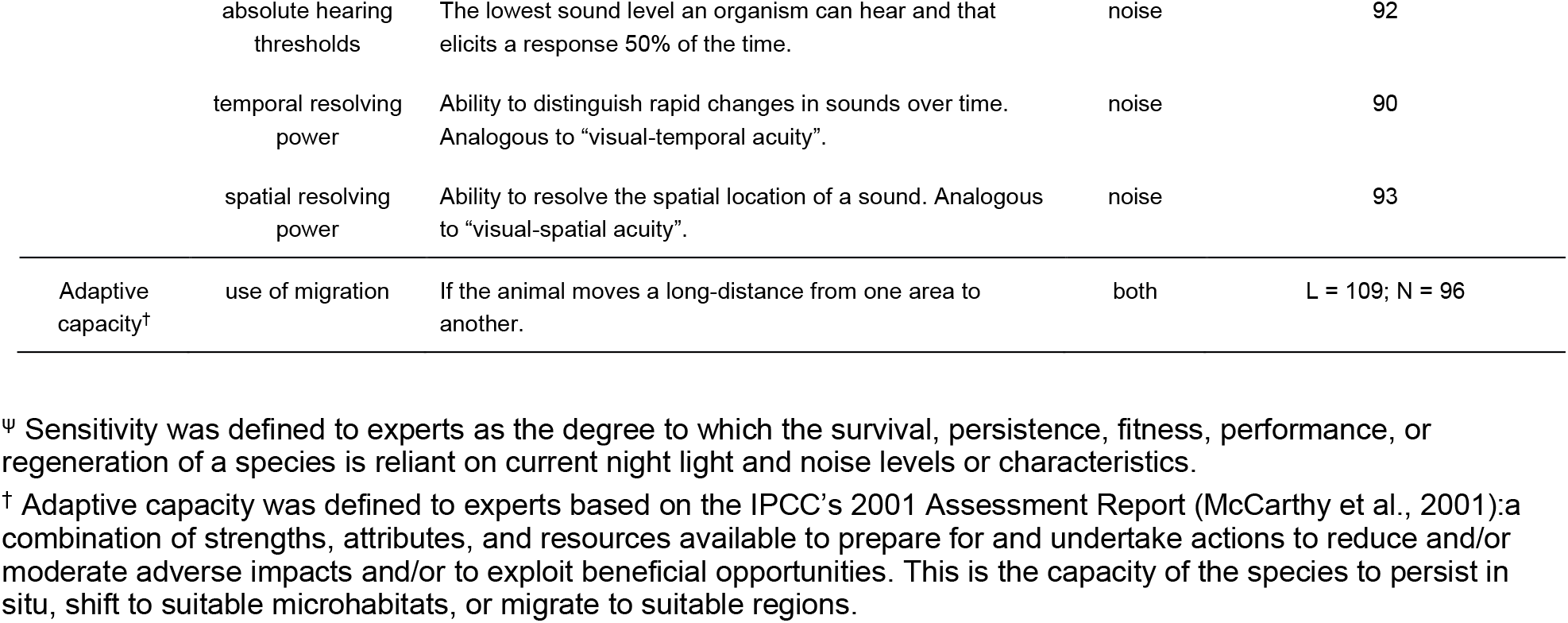
Summary of expert survey. Attributes include all biological traits that were assessed for their ability to increase vulnerability to anthropogenic night light and noise, the specific sensory pollutant we asked the expert to consider (light, noise or both) and the trait definition provided to expert within the survey. Experts were asked to assess the traits as they applied to all vertebrate species.

The survey began with two questions on the importance of anthropogenic light and noise within the systems the experts study and/or manage. The five response options to these questions ranged from “very important”, “important”, “moderately important”, “slightly important”, and “not important”. The next section elicited responses on specific traits and how each is related to vulnerability from lighting and noise. We asked experts about the impacts of lighting and noise (separately) on the same eight ecological sensitivity traits and “use of migration”. Because physiological and anatomical sensitivity traits were specific to either vision or hearing, we asked experts about the influence of lighting on six traits that were different from the seven traits considering the impacts of noise (**Table 1**).

We asked experts to provide a numeric value in response to three questions for each combination of sensory pollutant and trait. The first question asked about the importance of the trait and its effect on vulnerability if levels of lighting or noise are elevated. The six possible responses were: “0 – no effect”, “1 – small”, “2”, “3 – medium”,”4”, and “5 – large”. The second question asked which direction of the trait’s magnitude (“0 = no effect”, “1 – lowest/smallest/least”, up to “5 – highest/greatest/most”), or for some traits, specific categories (e.g., “nocturnal”, “diurnal”, or “no effect”) would increase vulnerability to lighting and noise the most (**Table 1, Figs. 1–4**). We then asked experts to assess the level of certainty in their responses (“0 – none” up to “5 – high”; see **Figs. 1–4** for details).

**Figure 1.).**
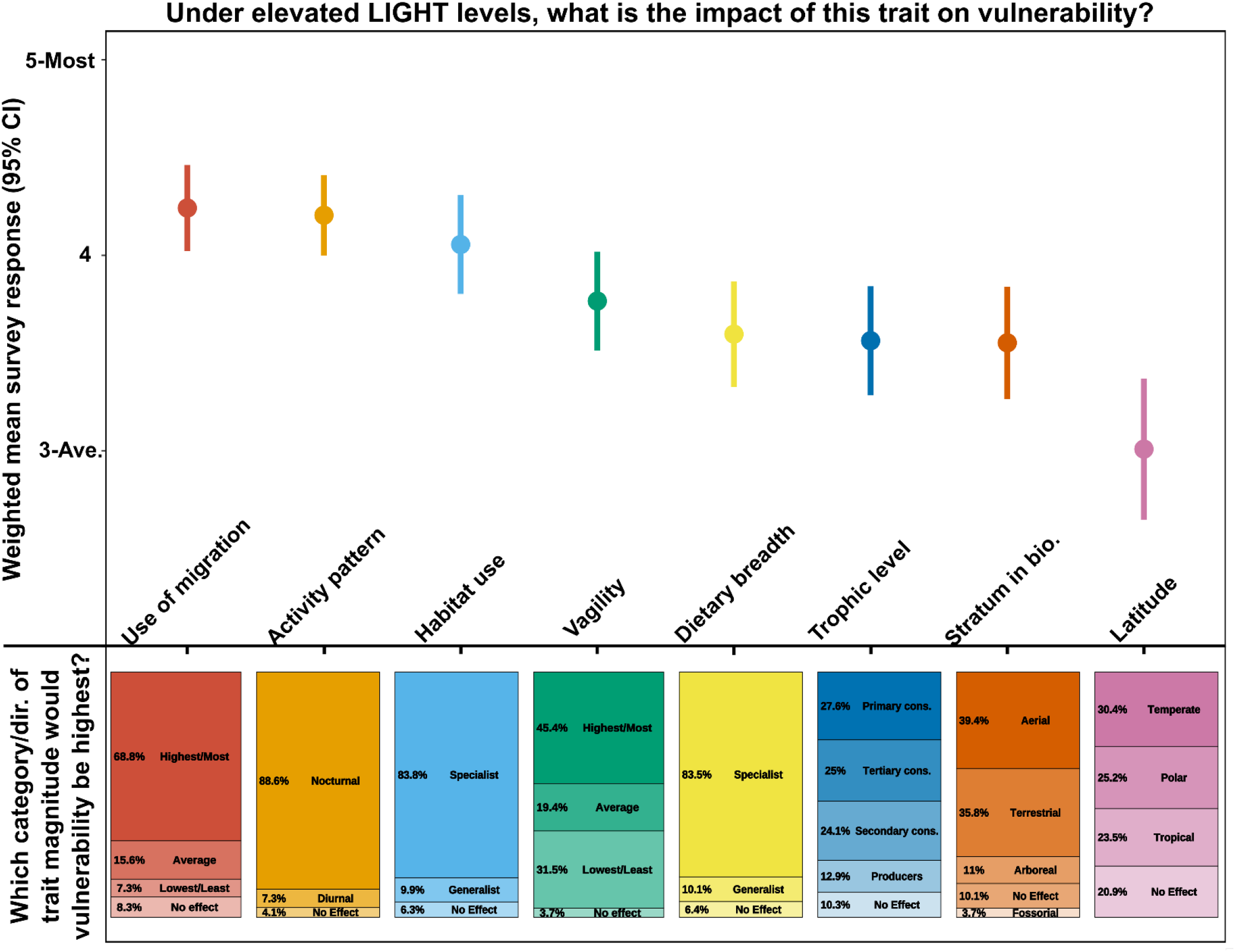
Results indicating the responses of experts assessing how ecological sensitivity and adaptive capacity traits influenced species’ vulnerability to anthropogenic night light, and whether having more/less of the trait, or specific attributes increased the magnitude of vulnerability. The 95% confidence intervals associated with the weighted mean vulnerability for each trait were derived using weighted errors from each respondent’s confidence in their answer. Confidence was scored [0 – 5 scale] as the following: 5 = “I have extensive knowledge of this trait and am very confident in my response”, 3 = “I have some knowledge of this trait and am moderately confident in my response”, 1 = “I have limited knowledge of this trait and am not confident in my response”, and 0 = “none”.

**Figure 2.).**
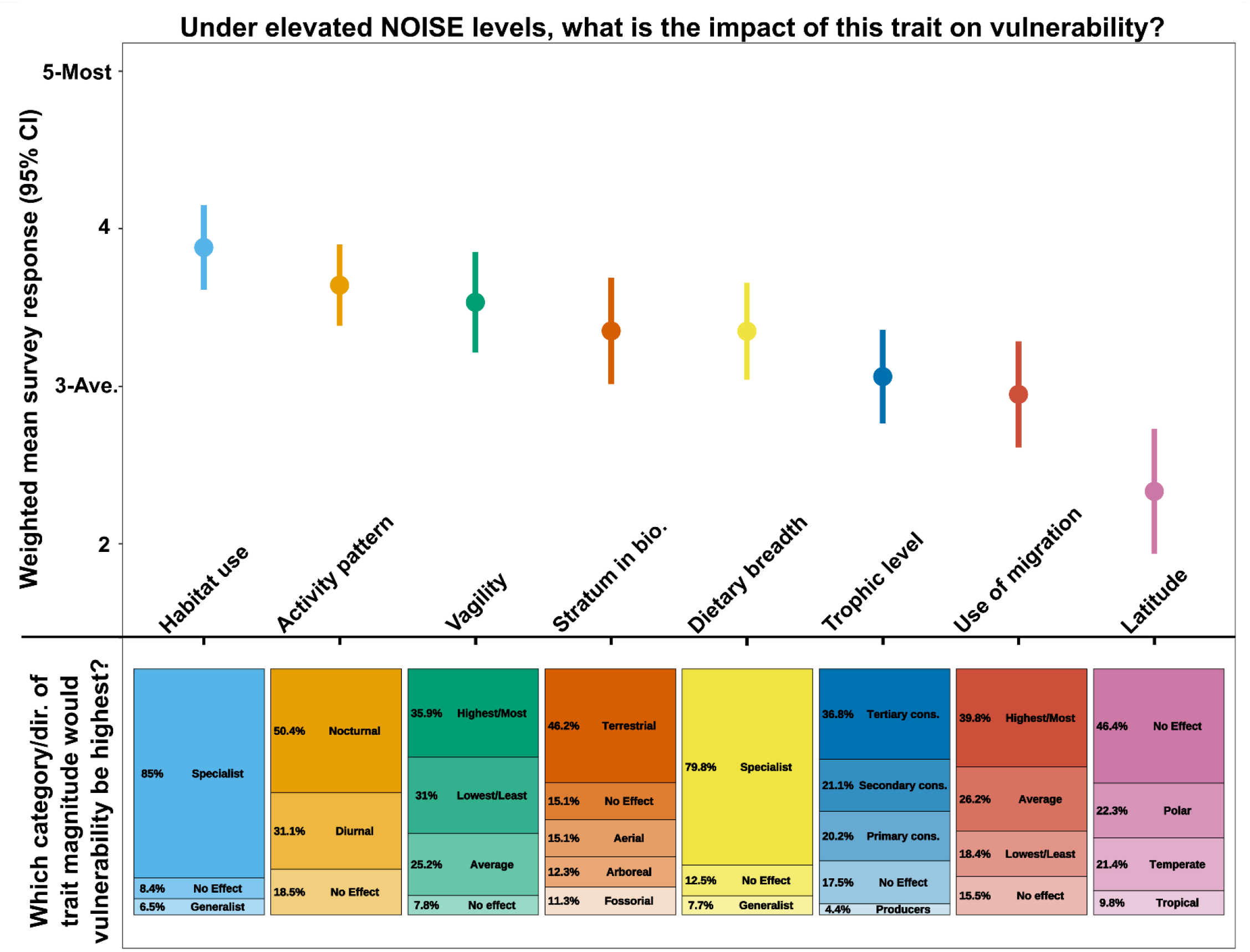
Results indicating the responses of experts assessing how ecological sensitivity and adaptive capacity traits influenced species’ vulnerability to anthropogenic noise, and whether having more/less of the trait, or specific attributes increased the magnitude of vulnerability. The 95% confidence intervals associated with the weighted mean vulnerability for each trait were derived using weighted errors from each respondent’s confidence in their answer. Confidence was scored [0 – 5 scale] as the following: 5 = “I have extensive knowledge of this trait and am very confident in my response”, 3 = “I have some knowledge of this trait and am moderately confident in my response”, 1 = “I have limited knowledge of this trait and am not confident in my response”, and 0 = “none”.

**Figure 3.).**
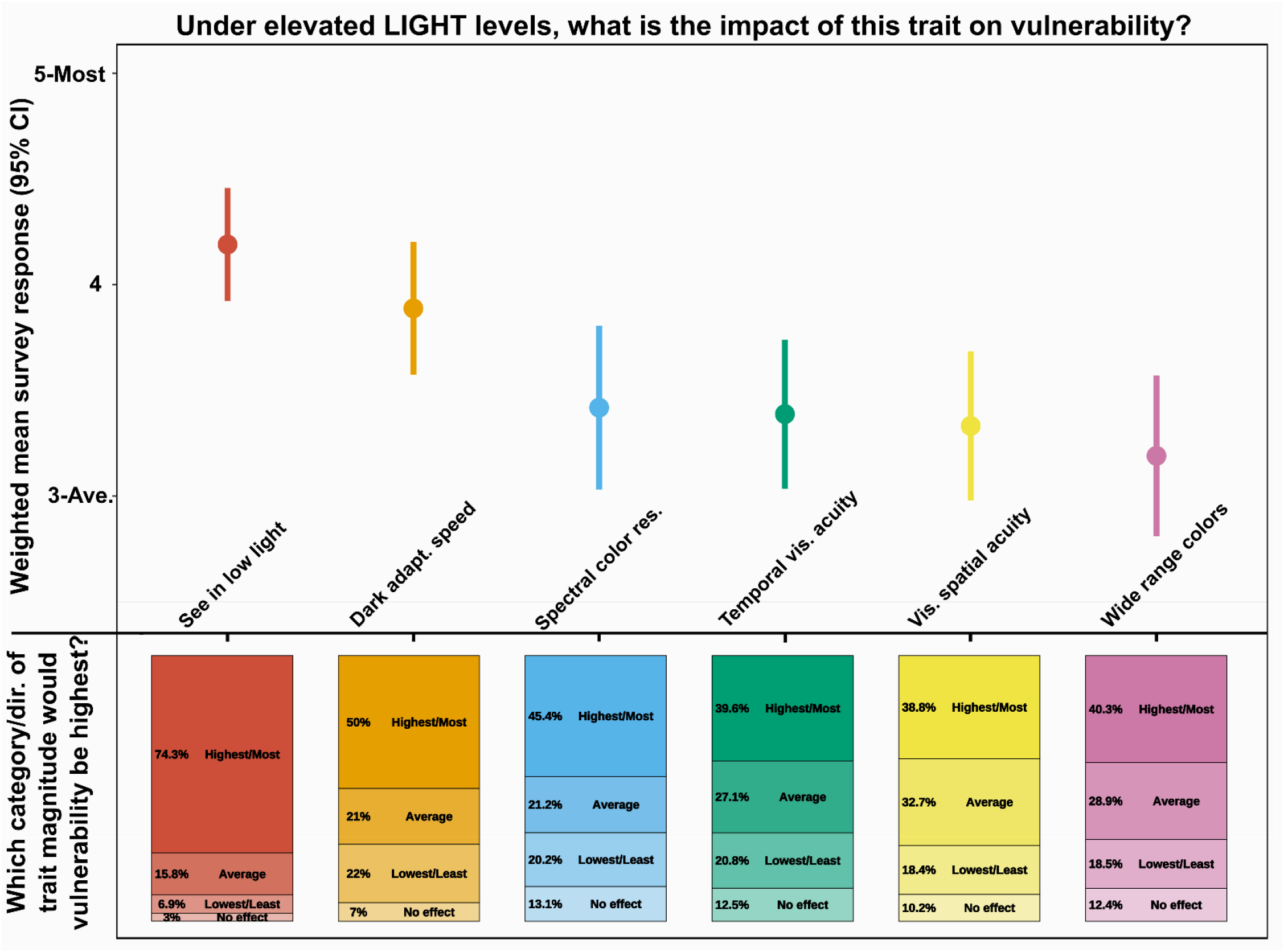
Results indicating the responses of experts assessing how anatomical and physiological sensitivity traits influenced species’ vulnerability to anthropogenic night light, and whether having more/less of the trait, or specific attributes increased the magnitude of vulnerability. The 95% confidence intervals associated with the weighted mean vulnerability for each trait were derived using weighted errors from each respondent’s confidence in their answer. Confidence was scored [0 – 5 scale] as the following: 5 = “I have extensive knowledge of this trait and am very confident in my response”, 3 = “I have some knowledge of this trait and am moderately confident in my response”, 1 = “I have limited knowledge of this trait and am not confident in my response”, and 0 = “none”.

**Figure 4.).**
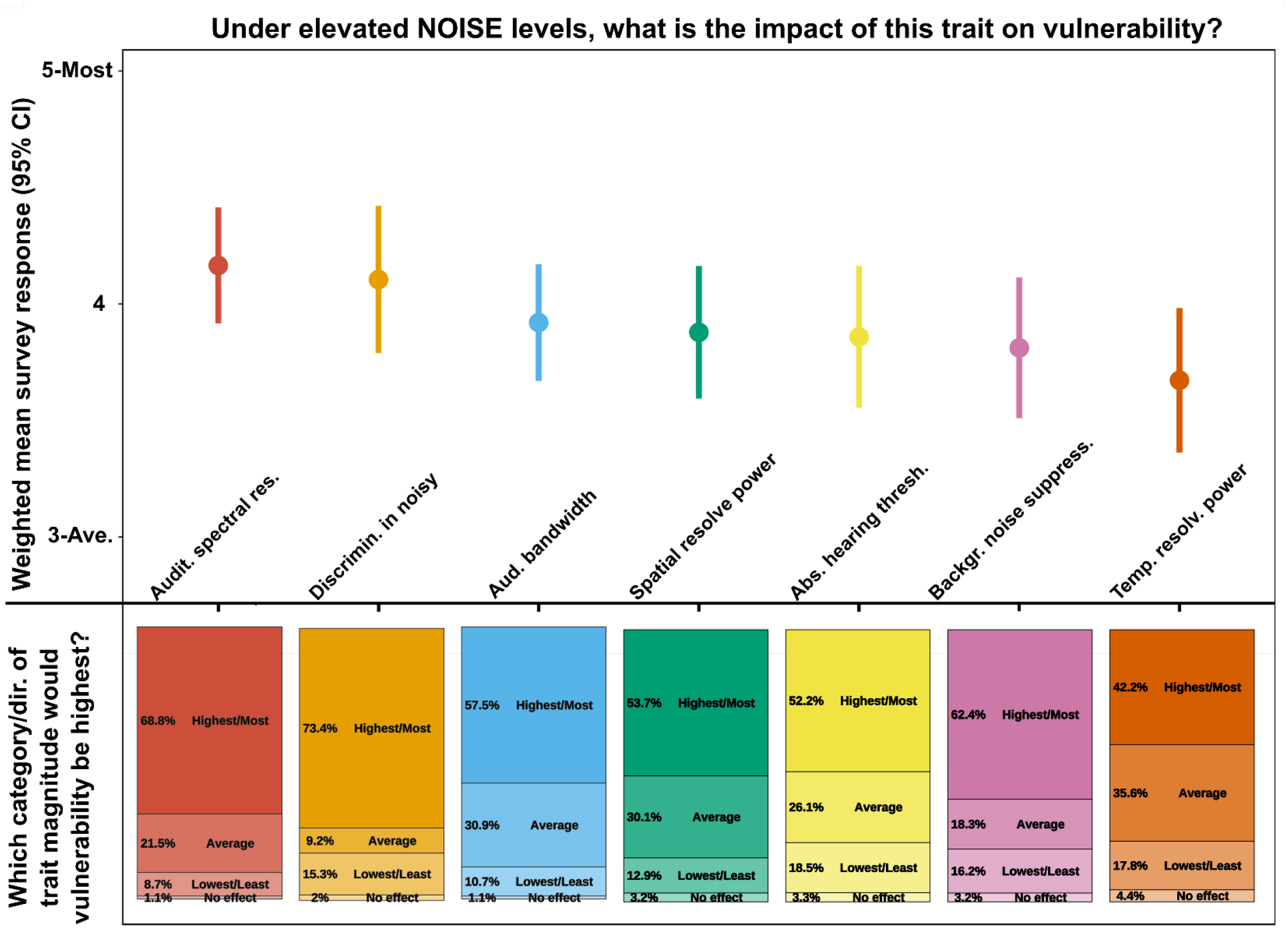
Results indicating the responses of experts assessing how anatomical and physiological sensitivity traits influenced species’ vulnerability to anthropogenic noise, and whether having more/less of the trait, or specific attributes increased the magnitude of vulnerability. The 95% confidence intervals associated with the weighted mean vulnerability for each trait were derived using weighted errors from each respondent’s confidence in their answer. Confidence was scored [0 – 5 scale] as the following: 5 = “I have extensive knowledge of this trait and am very confident in my response”, 3 = “I have some knowledge of this trait and am moderately confident in my response”, 1 = “I have limited knowledge of this trait and am not confident in my response”, and 0 = “none”.

### Survey data analysis

We calculated the weighted mean and weighted standard deviation of the responses indicating the level of influence on vulnerability to elevated levels of lighting or noise for each trait using the expert’s reported level of certainty to weight each metric. We used the package ‘diagis’ (Helske 2018) in program R (R Core Team 2019) to compute the estimates. The functions “weighted_mean” and “weighted_se” use probability weights instead of frequency weights. We constructed 95% confidence intervals by multiplying the weighted standard error by 1.96. The percentage of choices among options describing the direction of the trait’s magnitude were also calculated for each trait and sensory pollutant type.

### Survey Elicitation

We sought participants for our online survey from a variety of groups and organizations that regularly had contact with species’ experts. We first requested participation on popular email listservs, such as ECOLOG-L, and through snowball sampling in which survey participants recommend the survey to colleagues. We also searched Google Scholar for authors who were experts in wildlife and sensory ecology, producing a list of 135 and 34 potential participants, respectively. We sent emails to those authors requesting their participation in the survey.

When reaching out we stated that we were interested in participants that include, “sensory ecologists or those with ~3 or more years of study/experience with a particular vertebrate species/genus/taxon. A PhD candidate studying sea turtle nesting success, a biologist working in Everglades National Park for ten years, or an assistant professor would all potentially be suitable for this survey.” Respondents reported diverse areas of expertise, including “astrophysicist with experience in animal behavior”, and experience working with a variety of species (primary area of expertise: mammal(s) = 43%; bird(s) = 25%; amphibian(s) = 13%; invertebrate(s) = 8%; reptile(s) = 5%; fish = 5%).

Our survey provided an informed consent document to all participants that reminded the reader that participation was voluntary, it included detailed information on the purpose of the study, names and contacts of the principal investigators, and project sponsors. Participants were informed that we would make every effort to protect participants’ confidentiality and we asked participants to sign and date the informed consent form. The research protocols were approved by Boise State’s University Office of Research Compliance (approved IRB#: 193-SB18-068).

## Results

Nearly half of experts (48.4%; n = 280 responses) considered noise to be “very important” or “important” in the system each expert studies or manages, while 14.6% of experts considered noise “not important”. Slightly fewer experts (43.5%) considered lighting to be “very important” or “important”, and slightly more (20.0%) considered lighting to be “not important”.

### Vulnerability to lighting and noise based on ecological traits

Experts believed that elevated levels of lighting would increase the vulnerability of species that are highly migratory, are more nocturnally active, and are considered specialists when it comes to habitat use (**Figure 1**). Beyond these, several traits had similar, and lower, weighted mean survey responses. Although of moderate importance relative to other traits, there was consensus among the experts that dietary specialists have more vulnerability than dietary generalists to lighting and noise (**Figure 1**).

Habitat use specialists were considered most vulnerable to increased levels of noise (**Figure 2**). Activity pattern, vagility, stratum in the biosphere, and dietary breadth had similar weighted mean responses, but beyond dietary specialization, there was little consensus on the specific directionality or category of these traits (**Figure 2**).

### Vulnerability to lighting and noise based on anatomical and physiological traits

Respondents largely agreed (74.3%) that species with greater abilities to see in low light, and those with the fastest dark adaption speed (50% of responses) would be relatively more susceptible to light pollution (**Figure 3**). There was little difference among traits with lower weighted mean values but having the ability to see a wide range of colors (40.3% of responses) was considered to increase vulnerability the least (**Figure 3**).

When considering elevated levels of noise pollution, auditory spectral resolution and the ability to discriminate wavelengths of sound in noisy environments were considered most likely to increase the vulnerability of species (**Figure 4**). For both, experts generally agreed that having the most/highest ability of either trait increased vulnerability the most (68.8% and 73.4%, respectively; **Figure 4**). Temporal resolving power had the lowest mean survey response of all traits considered to influence the vulnerability to noise pollution (**Figure 4**).

## Discussion

We demonstrated that experts view sensory pollutants as important ecological stressors in the system they research or manage. Furthermore, the experts considered several specific traits, especially those related to having a narrow niche breadth, that make species more vulnerable to sensory pollution. These traits can serve as heuristics when considering disturbances from lighting and noise in developing policies for species that share the same traits.

In many ways, expert responses considering the threats from sensory pollutants aligned with assessments of climate change vulnerability (Julliard, Jiguet, & Couvet 2004; Clavel, Julliard, & Devictor 2011) by emphasizing the sensitivity of species with highly specialized habitat requirements. This analysis, however, expands upon this finding by focusing on the intersection of niche specialization with highly developed sensory function. The logic is simple: sensory degradation may critically depress productivity among habitat specialists. Although noise and lighting have not been featured in recovery plans for several habitat specialists, such as the spotted owl (*Strix occidentalis*; USFWS 2011), black-footed ferret (*Mustela nigripes*; see below & USFWS 2013), our analysis considering the input of hundreds of experts suggests they should be.

Here, we apply the framework of assessing sensory pollutant vulnerability to two endangered species to illustrate why noise and lighting management seems apt for their conservation plans. Gray bats (*Myotis grisescens*) have a nocturnal activity pattern, are habitat specialists (95% of the population roosts in 11 caves), and have eyes adapted to very low light levels. These traits, combined with the responsiveness of their prey to lighting, suggest they will be especially vulnerable to light pollution (**Figure 5**). Indeed, this species avoids areas affected by lighting (Cravens et al. 2018). Reduced light pollution can be realized by decreasing lumen output (or eliminating lights), better control over the spatial extent of lighting, limiting lighting to portions of the spectrum to which the bats and their prey are less sensitive, and limiting the seasonal and diel scheduling of lighting. For example, mitigating light pollution radiating from billboards or facades, and setting curfew hours for when they are turned on (Schroer et al. 2020), especially near the areas in which these animals roost, may be particularly effective.

**Figure 5.).**
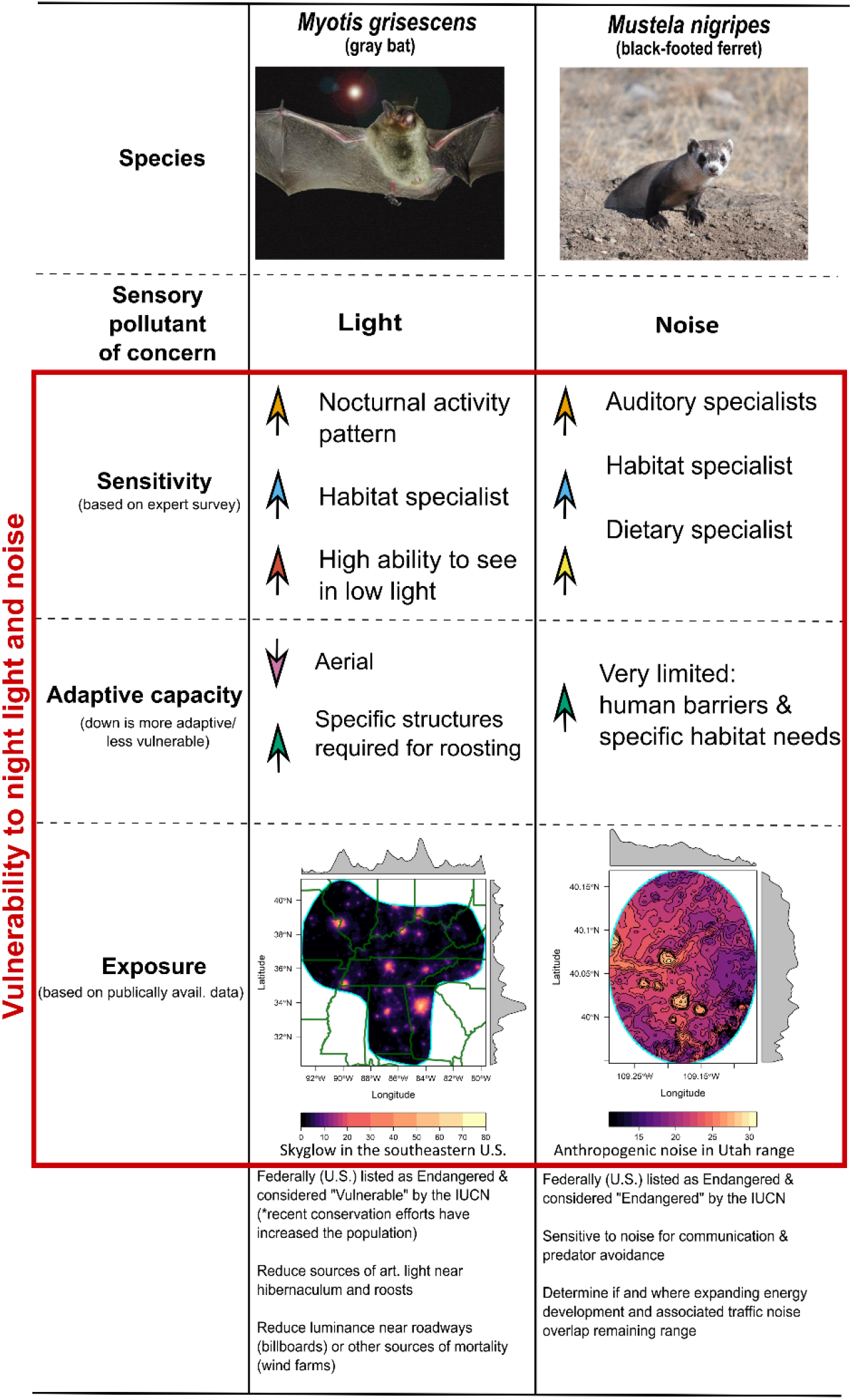
Applying expert survey to developing vulnerability risk assessments and potential future research and/or management actions. Here we selected two species of conservation concern to demonstrate how the results of our survey can be used to assess the overall vulnerability of a species to sensory pollutants. We used the entire species’ range of the gray bat in the southeastern U.S. and mapped the exposure based on estimates of nighttime skyglow developed by Duriscoe et al. 2018. For the black-footed ferret, we highlighted the species range in Utah only and mapped the estimated nighttime (ferrets are nocturnal) anthropogenic noise (L50) as developed by Mennitt & Fristrup (2016). Histograms show the mean values of each sensory pollutant across longitude and latitude.

The black-footed ferret has extremely specific habitat needs (USFWS 2013), and ferrets have exceptionally capable hearing on par with humans below 20kHz, and extending to an upper frequency limit of 40 kHz (Nodal and King 2014). Acute hearing is crucial for hunting in subterranean burrows and for avoiding predation by mesocarnivores above ground. Ferret ultrasonic hearing sensitivity enables them to eavesdrop on many rodent vocalizations that humans cannot hear. The prairie dog towns they require occur in open habitats lacking terrain shielding or sound attenuation due to vegetation, so noise propagates without obstruction. Noise can be reduced at the source through barriers, muffling, and scheduling of activities (Francis et al. 2011). For both of these endangered species, adaptive management could reduce these pollutants in a controlled experimental framework to quantify the benefits to these species and allow for mitigation methods to iteratively improve over time, while facilitating their adoption across many sites experiencing sensory pollution.

The aggregate responses of experts suggest that traits indicating highly developed sensory function — sensitivity to lower stimulus levels, better spectral resolution, better capacity to hear in noise or rapidly dark adapt after exposure to light — were generally regarded as indications that degraded sensory conditions would be more problematic. For the latter two traits, varied responses likely arose because some experts interpreted these traits as evolutionary evidence for heightened dependence on these senses, while others regarded these traits as evidence of better capacity to tolerate noise and lighting. Another grouping of responses exhibits similar divergence of responses. Stratum in the biosphere, vagility, and migration can be assessed from two perspectives. More vagile species may have more options to get away from adverse sensory conditions, mitigating the effects of these pollutants. Alternatively, more vagile species may be more heavily dependent upon sensory function for orientation, navigation, and surveillance in habitats where they have no recent experience. In the latter view, more philopatric species can use cognitive maps and recent familiarity with habitat conditions to offset some loss of sensory function. A dramatic demonstration of this latter perspective is the fatal, disorienting effects of light for highly migratory species (Van Doren et al. 2017; McLaren et al. 2018).

Diel activity patterns emerged as the second most emphasized ecological factor affecting sensitivity to noise and light (**Figs. 1, 2**). General consensus among experts points to sensitivity among nocturnal species that alter behavior in response to variation in artificial and natural light levels (Ditmer et al. 2020a, Willems et al. 2020, Prugh and Golden 2014), plus nocturnal acoustic specialists that respond negatively to noise exposure (Senzaki et al. 2016). Although the expert concordance was relatively high, it is possible that this general consensus may reflect sparse evidence among diurnal species, rather than an absence of effects. Studies of sensory function during sleep in wildlife are sparse; however, new studies are suggestive of impacts from both noise and light given the important role of hearing as a crucial alerting function during sleep and because light exposure appears to influence multiple physiological systems. Light disrupts the intensity, continuity and length of sleep in birds (Aulsebrook et al. 2020a,b) and noise appears to fragment and degrade sleep in birds much as it does in humans (Connelly et al. 2020). Thus, additional work is necessary to understand whether the costs of noise and light exposure are greater for nocturnal or diurnal species.

Notably, some of the expert responses differed from our expectations based on empirical studies. For example, we expected latitude to be considered an important trait for vulnerability to lighting (**Fig. 3**). In contrast, tropical populations have very consistent light cycles throughout the year, but changes in light radiance levels at twilight or loss of night could lead to misalignments in diel activity patterns within communities. In contrast, resident populations at high latitudes confront very long periods of night. When exposed to artificial light, the duration may be substantial. Tropical and temperate populations may also differ strongly in their responses to lighting depending on the degree to which variation in light regimes is a phenological cue. Many, but not all, temperate bird species appear to strongly advance their breeding season in response to lighting (Kempenaerns et al. 2010; Senzaki et al. 2020). For hearing, low absolute hearing thresholds would seem to be a prerequisite for elevated noise sensitivity. Some hearing experts might reasonably counter that critical ratios are the more important feature, but we were surprised by the lesser emphasis placed on this measure of auditory performance in our survey.

Although biological traits may change relatively slowly, lighting and noise are far less static, and may change dramatically within a population’s or a species’ range over the course of a single generation. Exposure is one of the three key components to vulnerability (Dawson et al. 2011). Geospatial models of skyglow have been developed (Falchi et al. 2016; Duriscoe et al. 2018), and Longcore et al. (2018) created an approach to predict species’ responses to spectral outputs based on behavioral and visual characteristics. Spatially explicit estimates of anthropogenic noise for the United States were developed by Mennitt & Fristrup (2016) and have been successfully applied to explaining how noise influences variation in avian reproductive success across North America (Senzaki et al. 2020). However, these geospatial models confront emerging challenges. The day-night band product from NASA’s VIIRS system cannot detect photons with wavelengths shorter than 500 nm. LED lamps that are rapidly proliferating through lighting upgrades have a prominent spectral peak at 470 nm, so global predictions of sky glow will require recalibration, and minimum estimates (Kyba et al. 2015a). The geospatial sound map was a composite created from ten years of measurements. More extensive monitoring and more sophisticated analyses will be required to produce the capacity to measure or predict trends.

The limitations of the study point to crucial future research directions. First, we only considered negative impacts from lighting and noise. However, future work should consider all effects, such as increased foraging opportunities for crepuscular species exposed to lighting (Santos et al. 2010) and enhanced ability to track resource peaks which are increasingly shifted temporally due to climate change (Senzaki et al. 2020). We also did not assess the impacts on invertebrates, an important group of animals that contributes a large percentage to many vertebrate diets, that are highly sensitive to changes in environmental cues (Klink et al. 2020; Owens et al. 2020), although 8% of respondents identified as experts of invertebrate species. In addition, our rankings only provide relative estimates of vulnerability. Combining these rankings with empirical measures of species response to sensory pollutants, such as reductions in survival, would mark an important advancement.

Despite the heightened understanding of the impacts that lighting and noise can pose to species, as demonstrated in our survey responses, and the increasing awareness that integrating sensory ecology is critical to conservation science (Dominoni et al. 2020), few conservation plans account for the expanding sensory footprint of the Anthropocene. Our study offers a generalized foundation for evaluating the ecological consequences of noise and lighting, and provides justification for management actions today. Although we understand enough to act now, and some governmental agencies are beginning to recognize the threats to wildlife from sensory pollutants and provide practical management solutions (see Mayer-Pinto, Dafforn, and Fobert 2020 for Australia), further studies are needed to determine the most economical and effective options to reduce sensory pollution at large enough scales to reduce harm to wildlife populations and ecosystem functions.

## Acknowledgements

The work was supported by the NASA Ecological Forecasting Grant NNX17AG36G. K. Markham assisted in developing and soliciting the survey.

